# Functional regulation of 4D metabolic network between multienzyme glucosome condensates and mitochondria

**DOI:** 10.1101/2022.05.16.491844

**Authors:** Erin L. Kennedy, Miji Jeon, Farhan Augustine, Krishna M. Chauhan, Songon An, Minjoung Kyoung

## Abstract

Glucose metabolism is biochemically intertwined between energy metabolism and building block biosynthesis in living cells. However, it has not been investigated yet how its metabolic network is orchestrated to govern glucose flux in space and time. Since we reported that human enzymes in glucose metabolism are spatially organized into metabolically active membraneless compartments (i.e., glucosomes), we have employed lattice light sheet microscopic imaging and other biophysical and biochemical techniques to understand their functional significance in cellular metabolism. Now, we demonstrated that glucosome assemblies behave like liquid droplets in human cells and thus reversibly respond to environmental changes. In addition, we characterized a molecular architecture of the glucosome, which appears to be constructed from higher-ordered oligomeric structures of its scaffolder enzyme along with transient enzyme-enzyme interactions. Importantly, we found that enzymatic compositions of glucosomes are altered when they are spatially in proximity to mitochondria to functionally couple glycolysis with mitochondrial metabolism in human cells. Collectively, we envision that the subcellular localization-function relationship between glucosomes and mitochondria may represent one of fundamental principles by which 4-dimensional metabolic networks are not only dynamically but also efficiently regulated in living human cells.

**One Sentence Summary:** Investigation of a 4D functional network of glucose metabolism uncovers a fundamental principal of subcellular metabolic regulation in human cells.

---

A metabolic network for energy production in cells mainly depends on a functional interplay between glycolysis and mitochondrial metabolism. Briefly, glycolysis breaks down glucose to pyruvate through a series of enzymatic activities in the cytoplasm. Pyruvate is then metabolized by mitochondria to create cellular energy via the tricarboxylic acid cycle and oxidative phosphorylation. In parallel, glycolytic intermediates can be shunted to building block biosynthesis such as the pentose phosphate pathway and serine biosynthesis for cellular biomass production. Such biochemical transformation of metabolic intermediates from glycolysis to mitochondrial metabolism as well as to building block biosynthesis has been well characterized at the biochemistry level. However, it has largely remained elusive how energy metabolism is selectively promoted at metabolic nodes over building block biosynthesis at subcellular levels in space and time.

Meanwhile, biomolecules such as proteins and/or nucleic acids have been identified to organize into biomolecular condensates spatially and temporally in cellular cytoplasm or nucleoplasm by liquid-liquid phase separation (LLPS)^1-5^. Such phase separated condensates are characterized as membraneless assemblies behaving like liquid droplets in living cells.

Importantly, biomolecular condensates have been functionally characterized to regulate specific biochemical reactions, respond to cellular environmental changes, localize specific molecules into defined subcellular locations, and/or modulate concentrations of biomolecules in cells^6^. In particular, we have discovered a multienzyme metabolic assembly, namely the “glucosome,” that encapsulates several enzymes participating in human glucose metabolism^7^. Subsequently, we have shown that glucosomes are spatially organized into various sizes to functionally orchestrate glucose flux between mitochondrial metabolism and building block biosynthesis^7,8^. However, it has not been elucidated yet how glucosomes are spatially regulated by LLPS to functionally communicate with other membraneless and/or membrane-bound organelles in living cells.

In this work, we employed various live-cell imaging platforms, particularly including lattice light sheet microscopic (LLSM) imaging, along with other biophysical and biochemical techniques, to quantitatively understand a 4-dimensional (4D) network of glucose metabolism in living human cells. We first visualized glucosomes using a fusion protein of human liver-type phosphofructokinase (PFKL) with a monomeric enhanced green fluorescent protein (i.e., PFKL-mEGFP) because it has been implicated as a glucosome marker in living human cells^7-9^. We then found that glucosomes are organized and regulated by LLPS in live cells. In addition, we revealed a molecular architecture of the glucosome that appear to be spatially constructed from higher-ordered oligomeric structures of PFKL and recruit other enzymes in glucose metabolism via transient enzyme-enzyme interactions. Importantly, we demonstrated that a functional network of glycolysis with mitochondrial metabolism is regulated by compositionally specific glucosomes in both proximity- and size-dependent manners. Collectively, we provide compelling evidence that multienzyme glucosome condensates are reversibly assembled near mitochondria by phase separation to selectively regulate glucose flux for energy metabolism in living human cells.

## Results

### Liquid-like phase behaviors of glucosomes in live Hs578T cells

Previously, we have demonstrated that a multienzyme glucosome assembly is organzied in the cytoplasm of human cells by enzymes that catalyze rate-determining steps in glycolysis or gluconeogenesis^7,9^, including but not limited to PFKL and muscle-type pyruvate kinase isoform 2 (PKM2). A fluorescnece recovery after photobleaching (FRAP) technique was then used to demonstrate that PFKL-mEGFP is capable of dynamically moving in and out of glucosomes^7^, revealing that glucosomes are membraneless compartments in human cells. However, their fusion and fission dynamics in living cells had not been definitivley characterized due to an intrinsically limited optical resolution on the z dimension of wide-field and confocal microscopic techniques. In this work, we employed time-lapse, sub-diffraction limited 3-dimensional (3D) LLSM imaging^10^ to address whether glucosome assemblies are regulated by LLPS in human cells^1,2,6,11^.

First, we investigated whether glucosome assemblies show liquid droplet-like behaviors in living cells. When we expressed PFKL-mEGFP as a glucosome marker in human mammary gland carcinoma cells (i.e., Hs578T), glucosomes were spatially displayed in various volumes and shapes in the cytoplasm of live Hs578T cells (Fig. 1a-d). Importantly, we found that glucosomes were temporally going through spontaneous fusion and fission processes under time-lapse 3D LLSM imaging (Fig. 1e., Fig. S1, and Movie S1), indicating they are pliable and undergo coalescent and pinch-off processes in live cells. To better understand such liquid-like behaviors of glucosomes, we analzyed an apparent concentration of PFKL-mEGFP in glucosomes by quantifying its fluorescence intensity per volumetric unit (i.e., voxel, (107 nm)^3^) from 3D LLSM images (e.g., Fig. 1e). We found that ratios of apparent concentrations of PFKL-mEGFP in glucosomes were statistically not changed (i.e., 0.94 ± 0.13) before and after fusion events. Alternatively, we quantified apparent concentrations of PFKL-mEGFP in randomly selected glucosomes from live Hs578T cells. Particularly when their volumes were ranged from 228.3 ± 20.3 to 969.3 ± 46.4 voxels, their apparent concentrations were no significantly different from each other (Fig. 1f, *n* = 2014). These data suggest that the fluorescence intensity and volume of PFKL-mEGFP in glucosomes are proportionally increased during fusion processes. In other words, apparent concentrations of PFKL-mEGFP in glucosomes remain constant or similar at any given moment between before and after a fusion event. Collectively, we reveal that glucosomes behave like liquid droplets while retaining their apparent concentrations during fusion and fisson processes in living cells.

**Fig. 1.**
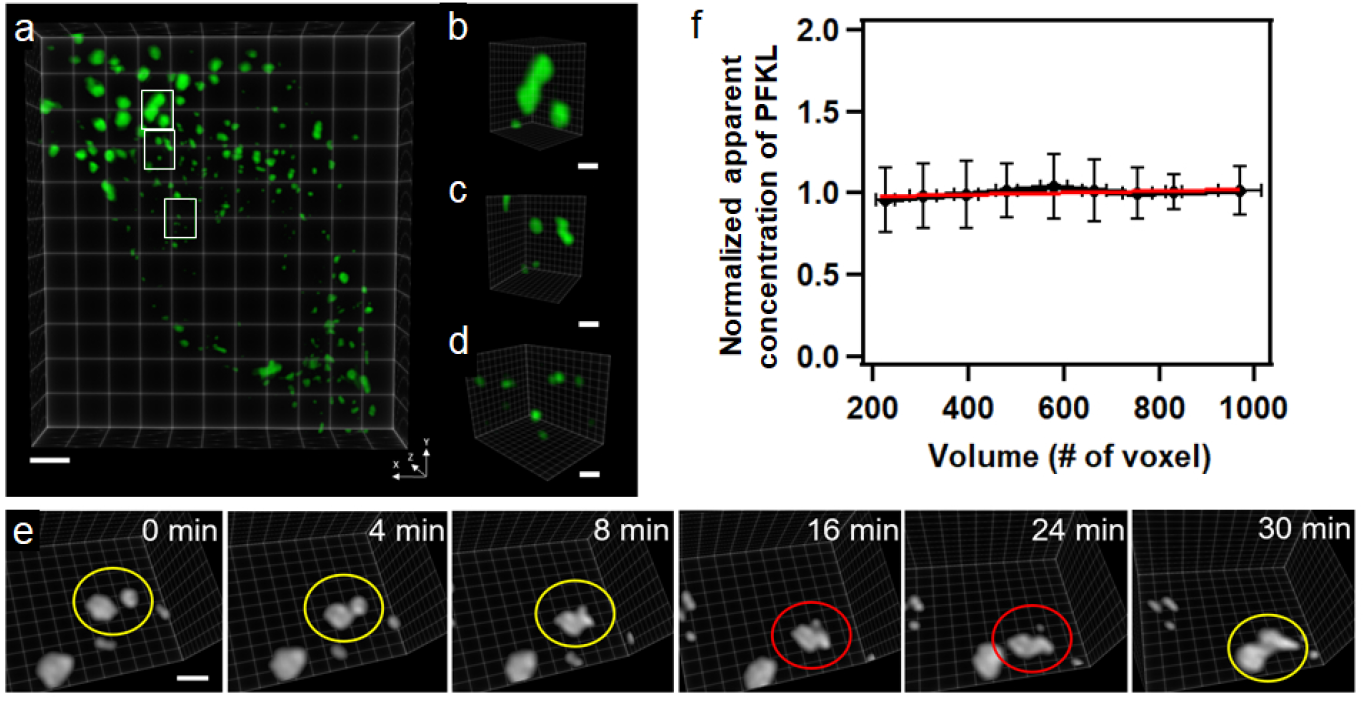
Liquid-like phase behaviors of glucosomes in live Hs578T cells. (a-d) Spatial distribution of PFKL-mEGFP as a glucosome marker in the cytoplasm of a representative Hs578T cell under 3D LLSM imaging. Scale bar in (a), 5 µm. Scale bars in (b), (c) and (d), 1 µm. (e) A selected set of image frames from a time-lapse imaging of a single live Hs578T cell showing fusion (yellow circles) and fission (red circles) processes of glucosomes. Scale bar, 5 µm. (f) Normalized apparent concentrations of PFKL-mEGFP in glucosomes as a function of glucosome volume (*N* = 2014). Error bars indicate standard deviations.

Second, we characterized whether glucosomes are regulated by LLPS during perturbation of celllular physicochemical properties. Of particular, we examined the effect of 4.4 % w/v polyethylene glycol 300 (PEG300) and 200 mM NaCl on glucosome reversibility in Hs578T cells because these are well-characterized to induce the same degree of hyperosmotic pressure on cell membranes^12,13^. Under wide-field fluorescence microscopy, we noticed that PFKL assemblies were significantly promoted in 72.4 ± 4.7 % of cells that were incubated with PEG300 for 20 min (Fig. 2a-b and Fig. 2d, red). When we subsequently removed PEG300, glucosomes were attenuated in 66.4 ± 5.3 % of cells (Fig. 2b-c and Fig 2d, green). Consistently, when a concentration of NaCl was increased from 135 mM to 200 mM, 78.8 ± 7.3 % of cells elevated the amount of glucosomes and subsequent reduction of NaCl to 135 mM reversely diminished glucosomes in cells (Fig. S2). At the same time, we subsequently analyzed 3D LLSM images in-depth before and after treatement of PEG300. We found that a total volume that glucosomes occupied per cell was drastically increased 249.2 ± 84.0 % by addition of PEG300 (Fig. 2e, red, *N* = 1834), relative to a no-treatment control, indicating that a dynamic partition of PFKL-mEGFP into its assemblies was significantly promoted. We also noticed that PEG300 slightly increased apparent concentrations of PFKL-mEGFP in glucosomes from 100 % to 112.3 ± 18.7 % under hyperosmotic pressure (Fig. 2f, *p* < 0.0001, *N* = 1834) while the volume of PEG300-treated cells was decreased about 10 % (Fig. S3). However, it is important to note that PEG300 treatment did not change an overall expresion level of PFKL-mEGFP per cell at single cell levels (Fig. S4). It appears that the observed reversibility of glucosomes under hyperosmotic pressure is mainly determined by the degree of dynamic partitioning of PFKL-mEGFP into glucosomes. Taken all together, we conclude that the reversibility of glucosome assemblies are certainly governed by LLPS in living human cells.

**Fig. 2.**
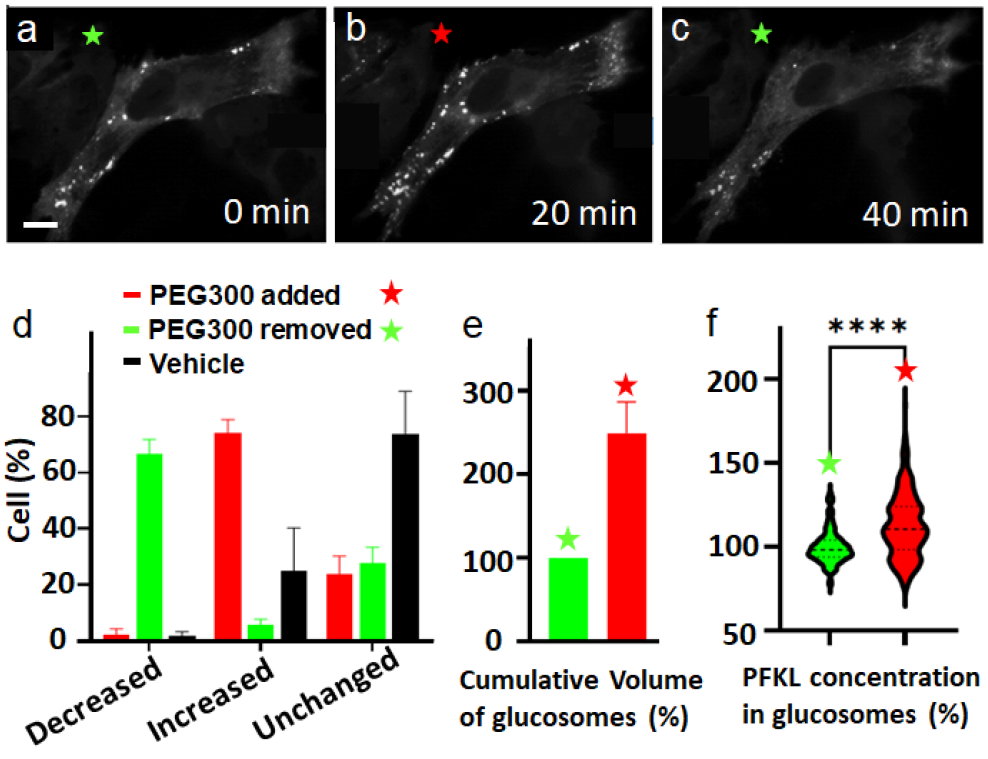
Glucosome reversibility under PEG300-triggered hyperosmotic pressure. (a-c) A representative Hs578T cell with glucosomes responding to addition and sequential removal of PEG300 (4.4 % w/v). (d) Percentages of cells showing decreased, increased, and unchanged amounts of glucosomes at single cell levels in response to PEG300 addition (red, *N*_*cells*_ = 83) and subsequent removal (green, *N*_*cells*_ = 83), along with a negative control using a vehicle (DMSO, black, *N*_*cells*_ = 78). Error bars indicate standard deviations from at least three independent experiments. (e) Cumulative volumes of glucosomes per cell in the presence (red) and absence (green) of PEG300 (*N*_*cells*_ = 9). Error bars indicate standard errors from at least 3 independent experiments. (f) Apparent concentrations of PFKL inside glucosomes in the presence (red) and absence (green) of PEG300 (*N*_*glucosomes*_ = 1834, **** *p* < 0.0001). At least 3 independent experiments were performed. Statistical analyses were performed using an unpaired t test.

### Molecular architecture of LLPS-promoted glucosome assemblies

We then investigated how enzyme-enzyme interactions within glucosomes contribute to the formation of LLPS-promoted glucosome assemblies. We first visulized colocalization of two enzymes in glucosome assemblies using mEGFP-tagged PKM2 (mEGFP-PKM2) and mCherry-fused PFKL (PFKL-mCherry) from live Hs578T cells (Fig. 3a). We then measured intracellular fluorescence resonance energy transfer (FRET) signals inside and outside glucosomes by quantifying an emisson intensity of a donor (i.e., mEGFP-PKM2) in response to an acceptor’s photobleaching as a function of time. We found that a FRET efficiency between PKM2 and PFKL was 11.5 ± 0.3 % (*N* = 33) on average inside glucosomes (Fig. 3b, green) whereas the donor’s emission outside glucosomes was comparable to the background fluorescence intensity (*N* = 15) (Fig. 3b, red). These results reveal that a direct interaction between PFKL and PKM2 is promoted inside glucosomes upon their colocalization in Hs578T cells. Next, to understand whether their interaction inside glucosomes are stable or transient in nature, we measured apparent diffusion coefficients of PFKL and PKM2 inside glucosomes using a FRAP technique. If a direct interaction between two enzymes would lead the formation of stable complexes within their assemblies, apparent diffusion coefficients (*D*) of both enzymes should be the same^14^. However, an apparent diffusion coefficient of PKM2 inside glucosomes (*D*_*PKM2,in*_) was 0.044 ± 0.022 µm^2^/s (*N* = 30), which was ∼ 2.4 times faster than a diffusion coefficient of PFKL inside glucosomes (*D*_*PFKL,in*_ = 0.019 ± 0.008 µm^2^/s, *N* = 51) (Fig. 3c), suggesting that they do not form a stable complex, and rather interact transiently inside glucosomes in Hs578T cells. Collectively, LLPS-mediated glucosomes appear to be assembled by direct but transient interactions among the enzymes in glucose metabolism.

**Fig. 3.**
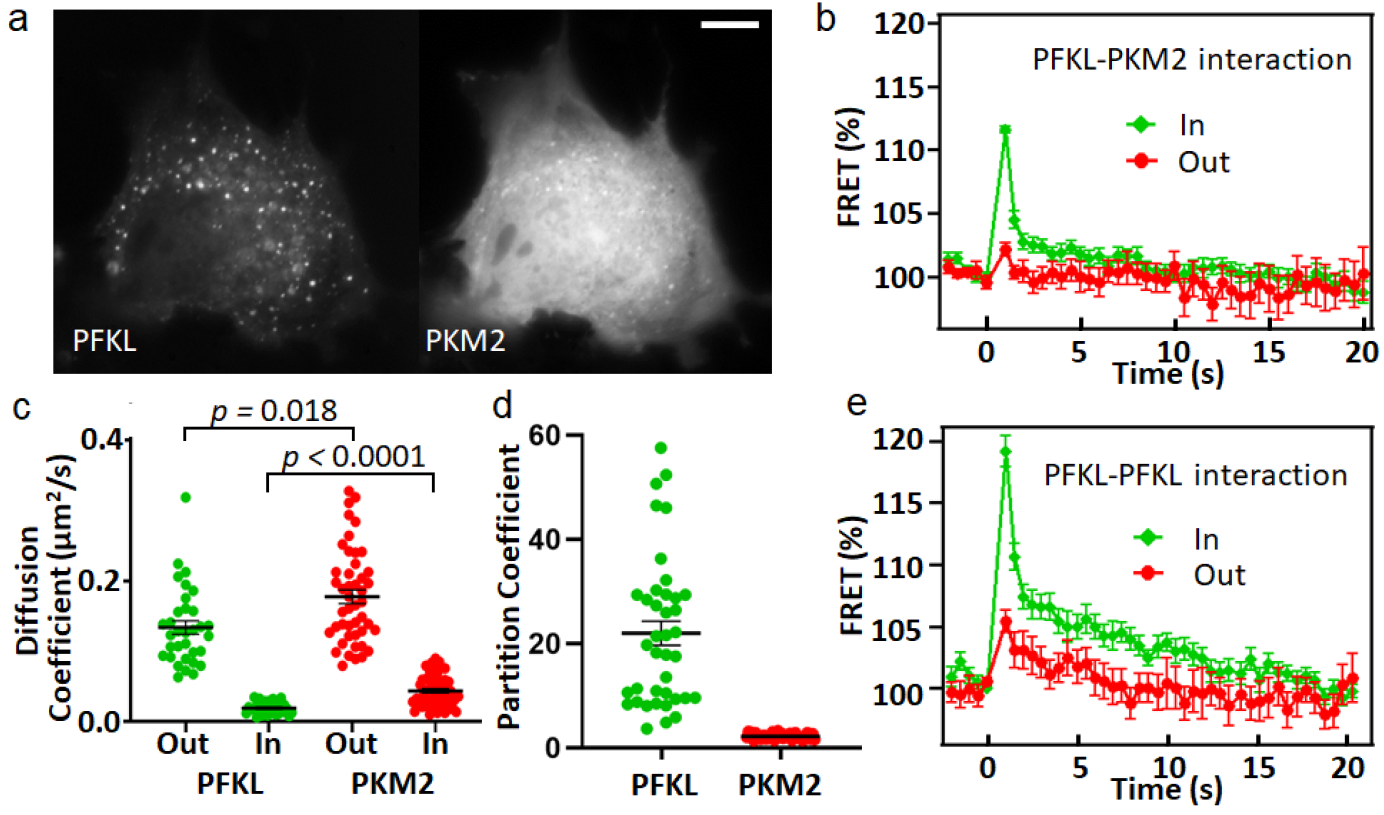
Molecular characteristics of PFKL and PKM2 inside and outside glucosomes in Hs578T cells. (a) A representative Hs578T cell showing colocalization of PFKL-mCherry and mEGFP-PKM2. Scale bars, 5 µm. (b) FRET efficiencies (%) obtained by an acceptor photobleaching method are shown for PFKL-PKM2 interactions inside (green, *N = 33*) and outside (red, *N = 15*) glucosomes. (c) Diffusion coefficients of PFKL-mCherry (green) and mEGFP-PKM2 (red) inside and outside glucosomes (*N*_*PKFL, out*_ *= 43, N*_*PKFL, in*_ *= 110, N*_*PKM2, out*_ *= 46*, and *N*_*PKM2, in*_ *= 45*). Error bars indicate standard errors. Statistical analyses were performed using an unpaired t test. (d) Partition coefficients of PFKL-mCherry (green, *N*_*PKFL*_ *= 39*) and mEGFP-PKM2 (red, *N*_*PKM2*_ *= 40*) from Hs578T cells. Error bars indicate standard errors. (e) FRET efficiencies (%) of PFKL-PFKL interactions inside (green, *N = 29*) and outside (red, *N =13*) glucosomes. Error bars indicate standard errors from at least 3 independent experiments.

In addition, we investigated whether PFKL would form higher-ordered oligomeric structures inside glucosomes to act as a scaffolder in living human cells. As far as we know, none of the pathway enzymes participating in glucosome assemblies^7^, including PFKL and PKM2, are yet characterized to form higher-ordered oligomeric structures in living cells. We first quantified fluorescence intensities of PFKL and PKM2 inside and outside glucosomes to analyze their apparent partition coefficients to glucosomes in Hs578T cells, respectively. We found that PFKL showed 22.02 ± 2.29 (*N* = 39) over PKM2 showing 2.24 ± 0.08 (*N* = 40) (Fig. 3d), indicating that PFKL partitions into glucosomes ∼10 times more than PKM2 does in our culture conditions. Subsequently, itimplies that PFKL may possess a certain molecular characetristic that would make it more favorable to participate in glucosomes than PKM2. We then compared ratios of apparent molecular weights (M.W.) of PFKL and PKM2 after their partition into glucosomes. In this analysis, apparent diffusion coefficients of PFKL and PKM2 inside glucosomes (Fig. 3c) were applied to the Stokes-Einstein equation with assumptions that both enzymes are spherical and have the same mass density^15^. Consequently, we obtained 12.4 as a calculated ratio of apparent M.W.s of PFKL-mCherry to mEGFP-PKM2 inside glucosomes. Since this is a significantly larger value than a sequence-based M.W. ratio (i.e., 1.3) of PFKL-mCherry (113.7 KDa) to mEGFP-PKM2 (85.7 KDa), such a high apparent M.W. of PFKL over PKM2 inside glucosomes may be explained if PFKLs form higher-ordered oligomeric structures, such as filaments as shown at least *in vitro* studies^16,17^, inside LLPS-promoted glucosomes.

To further examine whether PFKL forms oligomeric structures inside glucosomes, we analyzed intracellular FRET signals between PFKLs outside and inside glucosomes. We first co-expressed PFKL-mEGFP and PFKL-mCherry, in which each fluorescent protein was fused to the C-terminal of PFKL, in Hs578T cells. A FRET efficiency of 5.6 ± 1.0 % (*N* = 13) was measured between wild-type PFKLs *outside* glucosomes (Fig. 3e, red), resulting in 7.2 ± 0.2 nm for their average distance. This distance appears to be in good agreement with a distance of 7.5 nm that X-ray crystal structure of human platelet-type PFK (PFKP) (PDB: 4PFK) showed between the C-terminals of PFKP monomeric units within each dimer of its tetrameric structure^18^. We then, as a control, measured a FRET signal between PFKL F638R mutants (i.e., PFKL-F638R-mEGFP and PFKL-F638R-mCherry) in Hs578T cells because a F638R mutation was known to abolish a tetramerization of PFKL^17^. Their FRET efficiency of 4.9 ± 0.4 % (*N* = 48, Fig. S5) was indeed similar to what we observed between wild-type PFKLs outside glucosomes (i.e., 5.6 ± 1.0 %), supporting that two fluorescent proteins available at each dimeric unit of PFKL tetramers mainly contribute to the measured FRET signal between wild-type PFKLs outside glucosomes. Next, we measured an average FRET efficiency between wild-type PFKLs *inside* glucosomes, resulting in 19.2 ± 1.2 % (*N* = 29) (Fig. 3e, green). Accordingly, an average distance between the C-terminals of wild-type PFKLs was calculated as 5.5 ± 0.1 nm. This distance appears to be unique because it is shorter than any other distances we could measure between the C-terminals of PFKP monomeric units within a PFKP tetramer (i.e., 7.2 nm, 10.1 nm, or 10.0 nm). Our measured FRET-based distance between PFKLs inside glucosomes (i.e., 5.5 nm) appear to represent a distance between tetrameric units in a structure where they may be oligomerized. Taken all together with an apparent partition coefficient analysis and an apparent M.W. analysis of PFKL, our results here strongly support that PFKL forms higher-ordered oligomeric structures via enhanced PFKL-PFKL interactions inside glucosomes.

### Functional metabolic network between glucosomes and mitochondria

Meanwhile, we also investigated whether LLPS-promoted glucosomes are functionally coupled with mitochondrial metabolism in Hs578T cells. We designed three independent strategies to explore the functional metabolic network between glycolysis and mitochondrial metabolism in live Hs578T cells. First, we studied the effect of impairing oxidative phosphorylation in mitochondria using an inhibitor of mitochondrial ATP synthase (oligomycin A). Glucosomes and mitochondria were visualized with PFKL-mEGFP and MitoTracker-Red (Invitrogen) from live Hs578T cells using wide-field and LLSM imaging (e.g., Fig. 4a-c). When we then treated cells with oligomycin A (10 µM) for 1 h, 55.4 ± 7.1 % of Hs578T cells (*N* = 145) reduced the number of glucosomes under wide-field imaging at single cell levels (Fig. 4d). When we further analyzed those cells in detail under LLSM imaging, 37.1 ± 12.0 % of glucosomes per cell were impaired (Fig. S6). Apparently, glucosome assemblies were downregulated when oxidative phosphorylation was inhibited by oligomycin A in Hs578T cells. Second, we inhibited the pyruvate dehydrogenase complex (PDC) using a PDC inhibitor (CPI 613), that connects cytoplasmic glycolysis with the tricarboxylic acid cycle in mitochondria. In the presence of the PDC inhibitor (CPI 613, 500 nM), the decreased number of glucosomes per cell was observed in 82.3 ± 12.6 % of Hs578T cells under wide-field imaging (Fig. 4g, *N* =74).

**Fig. 4.**
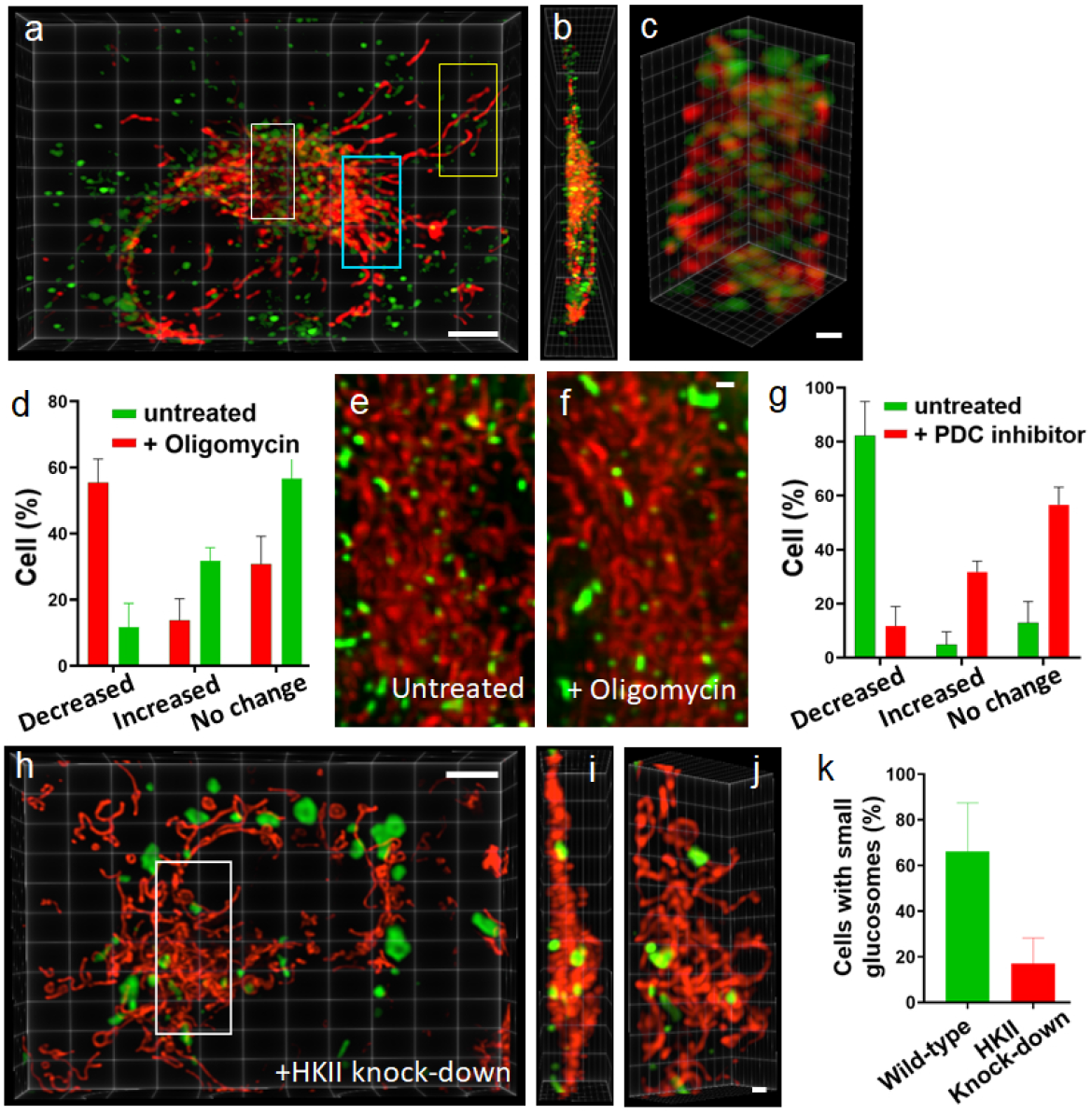
Metabolic association of glucosomes with mitochondria in Hs578T cells. (a-c) A representative Hs578T cell showing 3D distribution of glucosomes (PFKL-mEGFP, green) relative to mitochondria (MitoTracker-Red, red) under LLSM imaging. Scale bars in (a) and (c), 5 µm and 1 µm, respectively. (d) Percentages of cells showing decreased, increased, and unchanged amounts of glucosomes at single cell levels in the absence (green) and presence (red) of oligomycin A (10 µM) (*N*_*cells*_ = 145). Error bars indicate standard errors. (e-f) Representative cropped regions of interest in the cytoplasm of a live Hs578T cell showing relative spatial distribution of glucosomes (green) and mitochondria (red) in the absence (e) and presence (f) of oligomycin. Scale bars, 1 µm. (g) Percentages of cells showing decreased, increased, and unchanged amounts of glucosomes at single cell levels in the absence (green) and presence (red) of a PDC inhibitor ((6,8-Bis[(phenylmethyl)thio]octanoic acid (CPI 613), *N*_*cells*_ = 74). (h-j) A representative Hs578T cell showing spatial distributions of glucosomes (green) and mitochondria (red) after treatment of shRNA_HKII_. (i) and (j) show the side and the angled views of (h). Scale bars in (h) and (j), 5 µm and 1 µm, respectively. (k) Percentages of cells showing small-volume glucosomes with (red) and without (green) shRNA_HKII_ treatment. Error bars show standard errors from at least 3 independent experiments.

This observation was indeed consistent with the downregulation of glucosomes we observed in the presence of oligomycin A (Fig. 4d), supporting that there is indeed a functional metabolic link between membraneless glucosomes and membrane-bound mitochondria. Third, we downregulated glycolysis by knocking down hexokinase II (HKII) which enzymatically catalyzes the first and irreversible step of glycolysis. When Hs578T cells were transfected with small hairpin RNAs targeting HKII (shRNA_HKII_), western blot analysis confirmed that HKII was knocked down 52.8 ± 9.1 % in Hs578T cells (Fig. S7). When we subsequently applied shRNA_HKII_ under LLSM imaging, the number of small-volume glucosomes (< 90 voxels) per cell was noticeably diminished while the number of large-volume glucosomes (> 90 voxels) per cell was increased (Fig. 4h-j), relative to those observed from negative control cells without shRNA treatment (Fig. 4a-c). Quantitatively, shRNA_HKII_ treatment significantly decreased the population of cells showing small-volume glucosomes from 66.1 ± 21.4 % (Fig. 4k, green, *N =* 158) to 17.1 ± 11.2 % (Fig. 4k, red, *N* = 122). It is important to emphasize here that this study using shRNA_HKII_ not only corroborates that glucosomes are indeed functionally coupled with mitochondria, but also reveals that small-volume glucosomes may be responsible for the metabolic link between glycolysis and mitochondria. Collectively, we provide compelling evidence that functional decoupling between glycolysis and mitochondrial metabolism leads to a detrimental effect on glucosome assemblies, small-volume glucosomes, in live Hs578T cells.

### Spatial proximity representing a strong functional metabolic network between glucosomes and mitochondria

We then hypothesized that a functional link between glucosomes and mitochondria could be dependent on their spatial relationship in Hs578T cells. To test this hypothesis, we first analyzed, regardless of glucosome sizes, how close glucosomes were distributed relative to mitochondria in the cytoplasm of Hs578T cells under 3D LLSM imaging. To measure distances between mitochondria and glucosomes, we quantified a nearest edge-to-edge distance between two compartments from our collected 3D LLSM images (e.g., Fig. 5a, inset). We then found that, as shown in a histogram of the populations (%) of glucosomes per cell vs. their nearest edge-to-edge distance to a mitochondrion (Fig. 5a, *N* = 2147), a significant population (i.e., 37.8 ± 7.9 %) of glucosomes in single cells was indeed in closest proximity to mitochondria (i.e., the first bin of the histogram shown in Fig. 5a) within our 3D spatial resolution of LLSM (i.e., 275 nm). The histogram showing nearest edge-to-edge distances between glucosomes, and mitochondria was also well fit with a double exponential function but not with a single exponential function (Fig. 5a, solid black line), indicating there are two distinct subpopulations of glucosomes. They were apart from mitochondria in Hs578T cells with 65.4 ± 25.8 nm or 988.7 ± 41.9 nm based on decay distances of the double exponential fit. This data suggests that one subpopulation of glucosomes that are close to mitochondria may represent a strong spatial relationship with mitochondria whereas the other subpopulation of glucosomes appears to show stochastic distribution relative to mitochondria in Hs578T cells.

**Fig. 5.**
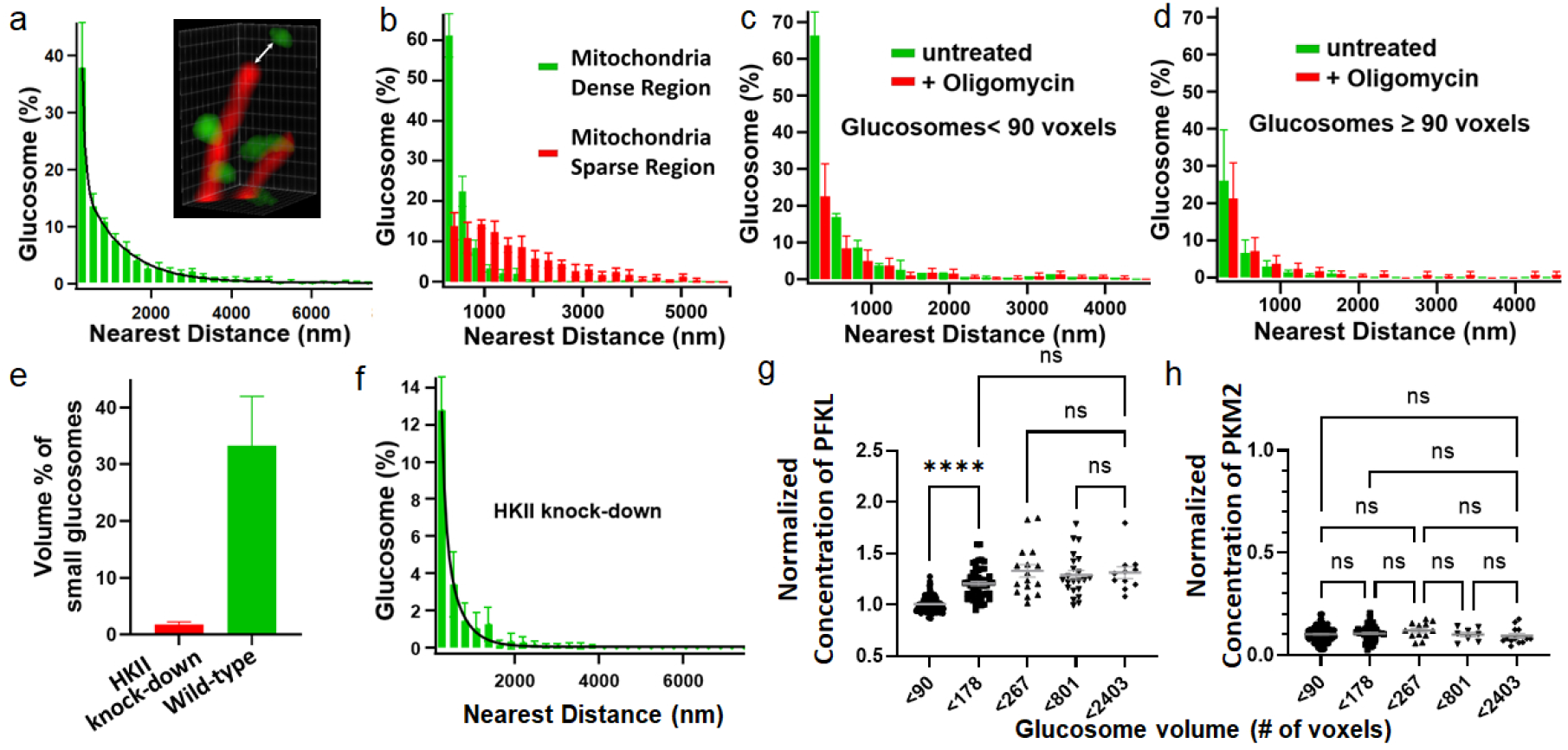
Spatial proximity-dependent functional link between glucosomes and mitochondria in Hs578T cells. (a) A histogram of glucosome populations (%) vs. the nearest edge-to-edge distance between glucosomes and mitochondria (*N*_*glucosomes*_ = 2147). A black solid line shows a double exponential fit for the histogram with a function (*f(d) = f(0) + A*_*1*_*e*^*-δ1·d*^ *+ A*_*2*_*e*^*-δ2·d*^) where mean distances for two subpopulations are expressed as *1/ δ1 and 1/ δ2*. A white arrow as shown in *inset* represents an example of the nearest edge-to-edge distance between a mitochondrion and a glucosome. (b) Histograms of glucosome populations (%) as a function of the nearest distance between glucosomes and mitochondria in mitochondria-dense (green) and mitochondria-sparse (red) regions (*N*_*glucosomes*_ = 899). (c-d) Histograms of populations of small-volume (< 90 voxels) and large-volume (≥ 90 voxels) glucosomes vs. the nearest distance between glucosomes and mitochondria only from mitochondria-dense regions in the presence (red) and absence (green) of oligomycin A treatment (*N*_*glucosomes*_ = 964). Error bars show standard errors. (e) Volume percentages (%) occupied by small-volume glucosomes at single cell levels with (red) (*N*_*cells*_ = 7) and without (green) shRNA_HKII_ treatment (*N*_*cells*_ = 8). (f) A histogram of glucosome populations (%) vs. the nearest distance between glucosomes and mitochondria after treatment of shRNA_HKII_ (*N*_*glucosomes*_ = 656). (g-h) Normalized apparent concentrations of PFKL and PKM2 as a function of glucosome volume. Note that the normalized apparent concentrations of PFKL and PKM2 are scaled based on the ratio of their partition coefficients (Fig. 3d). Statistical analyses were performed using a one-way analysis of variance (ANOVA). Statistical significance is defined as *p* < 0.05 with a 95 % confidence interval while ‘ns’ refers to not significant. * *p* < 0.05, ** *p* < 0.01, *** *p* < 0.001, **** *p* < 0.0001. Error bars indicate standard errors from at least 3 independent experiments.

During the edge-to-edge distance analysis, we also found that there were two types of distinct regions where mitochondria were densely packed or sparsely distributed in the cytoplasm of Hs578T cells. Subsequently, we analyzed spatial relationships of glucosomes with mitochondria at both regions under 3D LLSM imaging. In this analysis, we defined that a mitochondria-dense region was where mitochondria occupied on average 8.7 ± 1.3 % of a given space (e.g., Fig. 4a, blue square, Fig. S8a, and Movie S2) whereas a mitochondria-sparse region had mitochondria covering on average 1.8 ± 0.5 % of a given volume (e.g., Fig. 4a, yellow square, and Fig. S8b). Then, we found that in mitochondria-dense regions the majority of glucosomes (i.e., 61.2 ± 5.3 %, *N*_*Cells*_ *= 7*) were in closest proximity to mitochondria (Fig. 5b, green) while in mitochondria-sparse regions only 13.8 ± 3.4 % of glucosomes showed such a closest spatial relationship with mitochondria (Fig. 5b, red). Note that the number of glucosomes that were distributed in mitochondria-dense and mitochondria-sparse regions were statistically no different at least among our analyzed regions (Fig. S9, *p > 0*.*05*). Therefore, this analysis establishes that glucosomes that have a closer spatial relationship with mitochondria are mainly distributed in mitochondria-dense regions than in mitochondria-sparse regions.

Then, we examined what size of glucosomes were preferentially localized in mitochondria-dense regions and whether the degree of their spatial proximity to mitochondria is an important indicator for their functional communication. First, we measured volumes of glucosomes that showed a close spatial relationship with mitochondria in various mitochondria-dense regions. We found that 66.5 ± 12.1 % of glucosomes belonged to a category of small-volume glucosomes (i.e., < 90 voxels) under LLSM imaging. Second, we quantified changes of glucosomes that were localized in mitochondria-dense regions, regardless of their sizes, in response to treatment of oligomycin A under LLSM imaging. We observed that the population (%) of small-volume glucosomes that were the closest to mitochondria showed the most significant reduction from 63.5 ± 6.3 % to 22.5 ± 8.8 % with addition of oligomycin A (Fig. 5c). Then, as the distance from glucosomes to mitochondria increased, the degree of oligomycin-mediated reduction of small-volume glucosomes was significantly weakened (Fig. 5c). Conversely, larger glucosomes that had more than 89 voxels showed statistically no change with treatment of oligomycin A (Fig. 5d). These results support that the distance of small-volume glucosomes to mitochondria indeed represents the strength of their functional metabolic coupling in Hs578T cells. Third, we also analyzed subcellular volumes that were occupied by small-volume glucosomes in the absence and presence of shRNA^HKII^. Small-volume glucosomes in cells without shRNA_HKII_ was responsible for 33.3 ± 8.7 % of a total volume of all glucosome-occupied space in Hs578T cells (Fig. 5e, green). However, when cells were treated with shRNA_HKII_, the volume that was occupied by small-volume glucosomes was significantly reduced to 1.8 ± 0.5 % (Fig. 5e, red). Consistently, the population of glucosomes representing the closest spatial proximity to mitochondria (i.e., the first bin of each histogram shown in Fig. 5a & 5f) also showed significant decrease ∼3 times in the presence of shRNA_HKII_, relative to the corresponding population of glucosomes in no shRNA_HKII_-treated cells (i.e., 12.8 ± 1.8 %, Fig. 5f vs. 37.8 ± 7.9 %, Fig. 5a). These results support that downregulation of glycolysis with shRNA_HKII_ significantly decreased the population of small-volume glucosomes that were in close contact with mitochondria, thus decoupling its connection to mitochondria in Hs578T cells. Taken all together with our results showing a functional link between glucosomes and mitochondria (Fig. 4), our results strongly corroborate our conclusion that the closer glucosomes localize to mitochondria, the stronger their functional link with mitochondria is.

### Functional implication of glucosomes in enzyme stoichiometry-dependent manners

Since only small-volume glucosomes represented a strong spatial and -functional relationship with mitochondrial metabolism, we hypothesized that molecular characteristics of small-volume glucosomes may be distinctive to be functionally coupled with mitochondrial metabolism, relative to other sizes of glucosomes. To address this hypothesis, apparent concentrations of PFKL and PKM2 were analyzed as a function of glucosome volumes (Fig. 5g). Our analysis revealed that PFKL showed statistically no different concentrations in glucosomes when their volumes were in a range from 90 to 2404 voxels (Fig. 5g, *p* > 0.05). However, as the volume of glucosomes was less than 90 voxels (i.e., small-volume glucosomes), the concentration of PFKL decreased about 21.7% compared to its concentration in larger glucosomes (90 to 2404 voxels, *p* > 0.0001). In contrast, concentrations of PKM2 were statistically the same, regardless of volumes of glucosomes, including a case that their volume was < 90 voxels (Fig. 5h). It means that a compositional ratio of PKM2 to PFKL is higher in small-volume glucosomes (< 90 voxels) compared to their ratio in larger glucosomes (≥ 90 voxels). Therefore, this result suggests that small-volume glucosomes having a higher compositional ratio of PKM2 to PFKL may explain their strong functional commitment to glycolysis and thus their functional association with mitochondria.

## Discussion

Previously, we have demonstrated that enzymes in glucose metabolism are spatially organized to form multienzyme glucosome assemblies in living human cells^7-9^. Depending on their sizes, glucosomes are capable of regulating glucose flux between glycolysis and building block biosynthesis at single cell levels in human cells. However, it has remained elusive how glucosome assemblies are fundamentally organized in space and time and how such mechanisms ensure their functional contribution to glucose metabolism in cells. In this work, we used various imaging platforms, including wide-field, confocal and lattice light sheet microscopies, and associated biophysical techniques, like FRET and FRAP, to explore glucosomes and their functional network in living cells. Consequently, we provide compelling evidence that LLPS-promoted glucosomes are formed in various sizes with different enzymatic compositions, which dictate their metabolic function at subcellularly defined locations in human cells. We propose that the defined subcellular localization-function relationship between glucosomes and mitochondria may represent one of fundamental principles by which 4-dimensional metabolic networks are dynamically and efficiently regulated in living human cells.

Accordingly, we generated a ‘in-cell phase diagram’ (Fig. 6a) to explain dynamic nature of glucosomes in living human cells. Particularly, we demonstrated in this work that hyperosmotic pressure promoted glucosomes formation at cell population levels (Fig. 2d & S2). To describe this result, a phase boundary was plotted as an asymmetrical parabola shape with two variables, a pressure and a cellular concentration of PFKL. As a result, the overall shape of the phase boudary supports that as a pressure increases, a wider range of cellular concentrations of PFKL would favor glucosome formation inside the demixing regime (yellow area) (Fig. 6a).

**Fig. 6.**
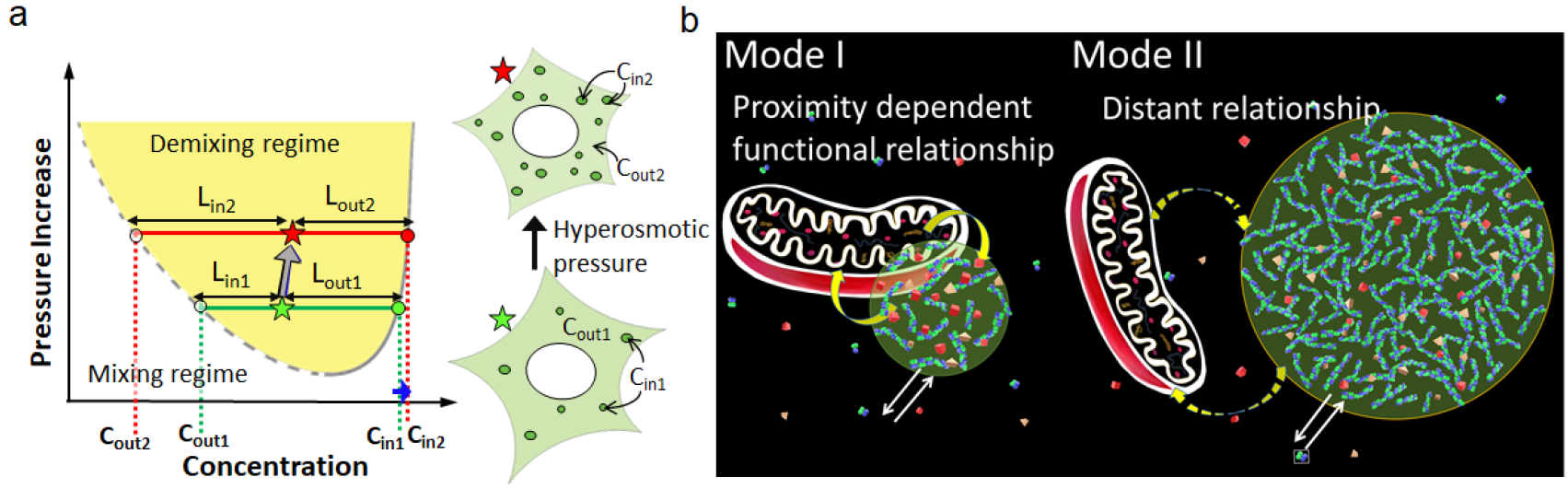
Schematic diagrams of LLPS-promoted glucosome in live cells. (a) An in-cell phase diagram describing glucosomes formation and their dynamics as a function of pressure (y-axis) and cellular concentration of PFKL (x-axis) in live cells. Green and Red stars represent a status of a single cell before (1) and after (2) osmotic pressure application. In tie-lines, L_x_ is a distance from a star-indicated location to the demixing boundary. C_in_ and C_out_ are intersections of a tie-line with phase boundary and also indicate a concentration of PFKL inside and outside glucosomes in a given cell. (b) Functional relationship between glucosomes and mitochondria in a size-dependent manner. Green spheres represent glucosomes. Cyan constituents forming glucosomes represent oligomeric PFKLs. Red and Brown constituents are client enzymes recruited by oligomeric PFKLs. Mode I depicts a strong functional link between a small-volume glucosome and a mitochondrion shown by solid curved arrows (yellow). Mode II depicts a weak functional link between a large-volumed glucosome and a mitochondrion by dashed curved arrows (yellow). White arrows present dynamic partitioning of enzymes into glucosomes.

We then enriched our phase diagram with more experimental data. As shown in Fig. 2e, a volume fraction of glucosomes was significantly increased as a pressure was raised at single cell levels. To accommodate this result, we introduce two tie-lines (i.e., green and red horizontal lines in Fig. 6a) in the demixing regime. All points at a green tie-line are assumed to be at ambient pressure, and the x coordinate of a green star indicates a cell having a specific cellular concentration of PFKL. Then, a volume fraction of PFKL in glucosomes is defined by the lever rule^6^, L_in1_/(L_in1_+L_out1_), where L_in1_ and L_out1_ are distances from a green star to two tie-line intersections with the phase bounday. At the same time, the increasing osmotic pressure is illustrated by a green tie-line translating to a red tie-line. Now, the x coordinate of a red star becomes slightly higher compared to that of a green star because our data demonstrated a slight increase of an apparent cellular concentration of PFKL under hyperosmotic pressure (Fig. S3-S4). We also found that a concentration of PFKL inside glucosomes (shown as C_in_ in Fig 6a) was only slightly elevated during the increasing osmotic pressure (Fig. 2f) (i.e., C_in1_ to C_in2_). Accordingly, the slope of the phase boundary at the high concentration range (solid grey, Fig. 6a) is depicted to be drastically steeper compared to its slope at the low concentration range (dashed grey, Fig. 6a). As a result, the increased osmotic pressure significantly raises a volume fraction of PFKL that forms glucosomes as shown from L_in1_/(L_in1_+L_out1_) to L_in2_/(L_in2_+L_out2_). Collectively, the illustrated in-cell phase diagram supports that human cells adapt to an increased pressure by preferably escalating PFKL participation into glucosome assemblies rather than significantly changing PFKL concentrations inside glucosomes in order to regulate glucosomes and thus glucose metabolism.

In addition, the subcellular architecture of a glucosome that we revealed in this work appear to be in a good agreement that biomolecular condensates are being formed particularly through multivalent interactions between hub molecules (i.e. scaffolders) and client molecules. Multivalent molecules are known to participate in biomolecular condensates with higher affinities than the affinities monovalent client molecules present^19,20^. Also, diffusions of multivalent molecules are anticipated to be slower than those of monovalent molecules due to their higher probablity of intermolecular interactions during diffusion and thus potentially heavier molecular weight after interactions. Accordingly, the higher partition coefficient (Fig. 3d) and the slower diffusion coefficient (Fig. 3c) of PFKL over PKM2 strongly suggest that PFKL may serve as a multivalent structural hub for glucosome condensates while PKM2 being a client. Moreover, intracellular FRET experiments (Fig. 3e) revealed higher-ordered oligomeric structures being formed by PFKL in live cells. Collectively, our results strongly suggest that oligomerized PFKLs may form a molecular architecture (Fig. 6b) that serves as mutivalent structural hubs for multienzyme glucosome formation and their dynamics in live cells.

Also, our FRET results demonstrating specific enzyme-enzyme interactions (Fig. 3b) suggest a potential mechnaism by which glucosome condensates are specific to the enzymes that are involved in only glucose metabolism. To date, many metabolic enzymes have been visualized to form various mesocale structures in living cells or *in vitro*^8,21-25^. Particularly, enzymes in *de novo* purine biosynthesis have been demonstrated to form spatially resolved and functionally active multienzyme assemblies in human cells (i.e., purinosomes^24^). However, when we visualized both glucosomes and purinosomes in single cells under 3D LLSM imaging (Fig.S10), it was apparent that they were spatially independent from each other in human cells. This strongly indicates that there is no possibilty of direct cross-pathway interactions through enzymes at least in our cell culture conditions. However, as we demonstrated in this work, specific enzyme-enzyme interactions occurs inside glucosomes (Fig. 3b), as well as insidepurinosomes ^14,26^, among their own pathway enzymes. In conjunction with our results revealing a subcellular molecular architecture of PFKL in glucosomes, we propose that compositional and thus metabolic specificity of glucosomes may be accomplished by having multivalent oligomeric PFKLs as structural hubs, followed by recruiting the other pathway enzymes only through specfiic interactions.

Importantly, we demonstrated dynamic spatial and functional relationships of glucosomes with mitochondria in live cells. Previously, we have determined that glucosomes are being organized in various sizes to regulate glucose flux in human cells. Particularly, small-sized glucosomes, which we defined to have < 0.1 µm^2^ based on an optical diffraction limit of mEGFP under wide-field imaging^7^, are proposed to regulate glycolytic flux over building block biosynthesis mainly based on our mathematical modeling study^27^. Now, we provide compelling experimental evidence that small-volume glucosomes under 3D LLSM imaging are indeed functionally responsible for regulating glycolysis for mitochondrial metabolism. Additionally, we demonstrated that small-volume glucosomes subcellularly localize near mitochondria through LLPS with a unique enzymatic composition to functionally control glycolytic flux for mitochondria. Collectively, it appears that a compositional ratio of participating enzymes, for instance a ratio of PKM2 to PFKL (Fig. 5g), within glucosomes determine how much glucose flux partitions between glycolysis and building block biosynthesis in a subcellular location-dependent manner.

In conclusion, we reveal a 4D mechanism of how glycolysis-promoting multienzyme glucosome condensates functionally communicate with mitochondrial metabolism in their proximity- and composition-dependent manners in living cells. We envision that this represents one of fundamental principles by which 4D metabolic networks, beyond glucose metabolism, are not only dynamically but also efficiently regulated in living human cells.

## Materials and Methods

### Materials

Plasmids expressing human liver-type phosphofructokinase with an enhanced green or mCherry fluorescent protein (PFKL-mEGFP or PFKL-mCherry, respectively), human muscle-type pyruvate kinase isoform 2 with mEGFP (mEGFP-PKM2), and human formylglycinamidine ribonucleotide synthase with mEGFP (FGAMS-mEGFP) were previously prepared^7,24^. MitoTracker™ Red CMXRos (Invitrogen, Cat# M7512) or MitoTracker® Green FM (Invitrogen, Cat# M7514) was used to stain mitochondria. Oligomycin A (Enzo Life Sciences, Cat# BML-CM111-0005) was used at a final working concentration of 10 µM. 6,8-Bis[(phenylmethyl)thio]octanoic acid (CPI 613, Tocris Bioscience, Cat# 5348), a PDH and KGDH inhibitor, was used at a final working concentration of 500 nM. Polyethylene glycol (PEG 300) and dimethyl sulfoxide (DMSO) were purchased from Sigma-Aldrich (Cat# 91462 and Cat# 276855, respectively). Blue/green/orange/dark-red TetraSpekTM microspheres were purchased from Invitrogen (Cat# T7280).

### Cell culture and transfection

Human breast cancer Hs578T cell line was purchased from the ATCC. Cells were maintained as previously described^7,9^. To prepare cells for transfection and subsequent imaging with wide-field and confocal microscopes, Hs578T cells were gently removed from a culture flask by trypsin-EDTA (Corning, Cat# 25-053-Cl) and seeded either glass-bottomed 35-mm Petri dishes (MatTek) or 8-well chambers (LabTek). For imaging with lattice light sheet microscope (LLSM), multiple 5 mm round coverslips (Warner Instruments, Cat# 64-0700) were placed on a 35 mm Petri dish (MatTEK) prior to seeding cells^9^. On the following day, cells were transfected with Lipofectamine 2000 (Invitrogen) using Opti-MEM-I reduced serum medium (Opti-MEM-I; Gibco, Cat# 11058). The transfection medium was then exchanged with fresh antibiotic-free growth medium (i.e., RPMI 1640 (Mediatech, Cat# 10-040-CV) and 10 % dialyzed fetal bovine serum (FBS, Atlanta Biological, Cat# S12850)) after a 5-h incubation (37 °C, 5 % CO_2_, and 95 % humidity), followed by ∼18–24 h of incubation in the CO_2_ incubator.

### Human hexokinase II (HKII) knockdown by short-hairpin RNAs (shRNAs)

Lentiviral pGFP-shHKII vector encoding human shRNA_HKII_ (Cat# TL312415) and non-silencing control pGFP-C-shLenti vector (Cat#: TR30023) were purchased from OriGene Technologies Inc (Rockville, MD, USA). To knockdown HKII, Hs578T cells (1 × 10^6^) were cultured overnight in 30 mm dishes containing RPMI 1640 and 10 % dialyzed FBS without antibiotics. On a next day, 1 µg of shRNA-expressing plasmids were used to transfect cells with Lipofectamine 2000 (Thermo Fisher Scientific, USA). After 24 h of transfection, a fresh growth medium (i.e., RPMI 1640 and 10 % dialyzed FBS) containing 1 μg/ml puromycin (Sigma-Aldrich, Cat# P8833) was added and subsequently puromycin-resistant cells were selected after 48 h.

Then, cells were washed with cold 1xPBS and subsequently lysed with RIPA buffer (150 mM sodium chloride, 1.0 % Triton X-100, 0.5 % sodium deoxycholate, 0.1 % sodium dodecyl sulfate, 50 mM Tris, pH 8.0) containing protease (Pierce, Cat# 88666) and phosphatase inhibitors (Pierce, Cat# 88667). Cell lysates were then used for western blot analysis.

### Fluorescence live-cell imaging

Wide-field and confocal imaging were carried out as previously described^7,9^. Before imaging, cells were washed with buffered-saline solution (20 mm HEPES, pH 7.4, 135 mm NaCl, 5 mm KCl, 1 mm MgCl_2_, 1.8 mm CaCl_2_, and 5.6 mm glucose) for three times with 10-min incubations, followed by a ∼1–2-h incubation at ambient temperature. All samples were then imaged at ambient temperature (∼25 °C) with a 60× 1.45 NA objective lens (Nikon CFI Plan Apo TIRF) using a Photometrics CoolSnap EZ monochrome CCD camera on a Nikon Eclipse Ti inverted C2 confocal microscope. Wide-field imaging was carried out using the following filter sets from Chroma Technology: mEGFP detection by a set of Z488/10-HC clean-up, HC TIRF dichroic, and 525/50-HC emission filter; and mCherry detection by a set of Z561/10-HC clean-up, HC TIRF dichroic, and 600/50-HC emission filter.

### Fluorescence recovery after photobleaching (FRAP)

FRAP was performed with live Hs578T cells as described previously^7,9^. Briefly, confocal imaging was performed using a JDSU argon ion 488-nm laser line and a Coherent sapphire 561/20-nm laser line for mEGFP and mCherry fusion proteins, respectively. Fluorescent signals in subcellular regions of interest (1-3 µm in diameter) in live cells were photobleached.

Subsequently, fluorescence recoveries at the photobleached regions were quantified for at least 50 seconds and then fitted individually after the degree of background photobleaching was normalized. Apparent diffusion coefficients (*D*_app_) were then calculated as we described before^7,9^.

### Fluorescence resonance energy transfer (FRET)

FRET measurements in live cells were also performed as described previously^7,9^. First, mEGFP-tagged donor and mCherry-tagged acceptor enzymes were dually transfected into Hs578T cells. On the next day, to measure FRET signals between donors and acceptors inside and outside glucosomes, respectively, we applied a Coherent sapphire 561/20-nm laser to photobleach mCherry-tagged acceptors in regions of interest. At least 10 images were obtained before the acceptor photobleaching. Right after the acceptor photobleaching, images were acquired every 0.5 s for at least 50 s. Temporal increases of the emission of mEGFP-tagged donors from the same regions were monitored to measure positive FRET signals from live cells. Dual-color confocal imaging was achieved via a 488/561/640 dichroic mirror with 525/50 and 600/50 emission filters and photomultipliers. Distances between donors and acceptors were calculated by *E*_*FRET*_ *= 1/[1 + (r/R*_*0*_)^*6*^], where r is a distance between the FRET pair and R_0_ is the characteristic distance for the given FRET pair.

### Lattice light sheet microscopic (LLSM) imaging

3D imaging was performed with a home-built LLSM as previously described^9^. On the day of imaging (∼18-24 h post-transfection), 100-200 nm blue/green/orange/dark-red TetraSpek™ microspheres (Invitrogen, Cat# T7280) were added to cell dishes to verify color registration using an in-house Matlab code for sub-pixel corrections in 3D^9^. They were also used to determine a daily point spread function in relevant color channels for a deconvolution process as described in detail before^9^. Additionally, hex-or square-patterned phase masks on the spatial light modulator were used to create thin plane illumination with a sub-diffraction limited z-dimensional resolution. Cells were then imaged at ambient temperature (25°C) with a 25x 1.1 NA objective lens (Nikon, CFI Apo LWD) using a sCMOS camera (Hamamatsu, Orca Flash 4.0 v2 camera) with a quad-band pass filter (Semrock, Cat# FF01-446/523/600/677-25).

### LLSM Image analyses

#### 3D edge-to-edge distance analysis

Binary object maps of 3D LLSM images for each color channel were generated using the ImageJ processing software (National Institutes of Health, NIH) or the Allen Cell Structure Segmenter^28^. These binary segmented files were analyzed using an in-house Matlab code to determine a shortest distance in 3D between surfaces of objects in the red and green channels. Briefly, the Matlab code read two input binary object maps and labeled coordinates of all objects. 3-Dimensional perimeter of each object was then defined using the regionprops and bwperim functions from Matlab. Two objects from each color channel were analyzed in pair-wise manner to calculate a vector distance between objects’ perimeter voxels using the bwdist function. Calculated distances were recorded as edge-to-edge distances for a pair of given objects.

#### Analysis of apparent concentrations of enzymes

3D cell images were processed using a 3D objects counter plugin from the ImageJ processing software^29^. Physical characteristics such as a volume defined by the number of voxels and an integrated fluorescence intensity of an object were obtained from the 3D object counter. An apparent concentration of an enzyme inside glucosomes was then calculated by dividing the integrated fluorescent intensity by the volume.

#### Analysis of apparent molecular weight ratios

To calculate an apparent molecular weight ratio of two enzymes, we first deduced a radius (*R*_*e*_) of each enzyme using the Stokes-Einstein equation, 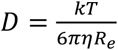 where k is the Boltzman constant, T is the temperature, η is the viscosity, and D is a diffusion coefficient measured from our FRAP experiments. Since this analysis is based on assumptions that an enzyme is spherically shaped and importantly that molecular densities of two enzymes are the same, a ratio of two enzymes’ apparent molecular weights was calculated from radii of PFKL and PKM2.

## Acknowledgments

A lattice light-sheet microscope was built in-house in Dr. Kyoung’s laboratory at UMBC after obtaining a research license agreement with the Howard Hughes Medical Institute (HHMI). We thank Eric Betzig (HHMI, Janelia Research Campus) and Wesley R. Legant (HHMI, Janelia Research Campus) for their generous support by providing technical helps and operational expertise. We also acknowledge Alicia D. An for drawing schematic illustrations. This work was funded by the National Institutes of Health; R01GM134086 (M.K.), R01GM125981 (S.A.), R25GM55036 (E.L.K.) and T32GM066706 (E.L.K. and M.J.). The content is solely the responsibility of the authors and does not necessarily represent the official views of the National Institutes of Health.

## Author Contributions

M.K. and S.A. designed the research, M.K. directed the research, E.L.K., M.J., F.A., K.M.C. and M.K. performed experiments and/or analyzed data, M.K. and S.A. wrote the manuscript. All authors reviewed and edited the manuscript.

### Data availability

All data are available within the main manuscript and the supplementary materials.

