## Supplemental Figures for "Functional regulation of 4D metabolic network between multienzyme glucosome condensates and mitochondria"

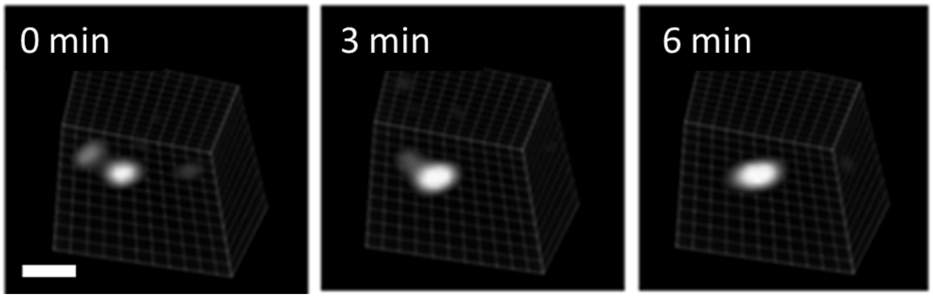


**Fig. S1. A selected set of image frames from a time-lapse imaging of a single live Hs578T cell showing a fusion event of glucosomes.** Scale bar, 0.5 μm.

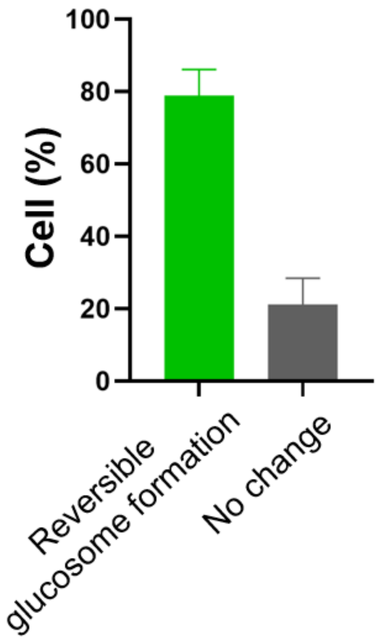


**Fig. S2. Glucosome reversibility under NaCl-triggered hyperosmotic pressure.** Percentage (%) of Hs578T cells was quantified when NaCl concentration was increased from 135 mM to 200 mM and subsequently decreased to 135 mM (N = 91). Error bars indicate standard error from at least 3 independent experiments.

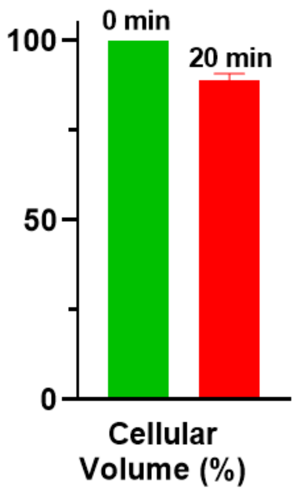


**Fig. S3. Effect of NaCl concentration on single-cell volume.** Volumes of single cells were quantified by 3D LLSM imaging when NaCl concentration was increased from 135 mM to 200 mM (N = 4). Error bars indicate standard error from at least 3 independent experiments.

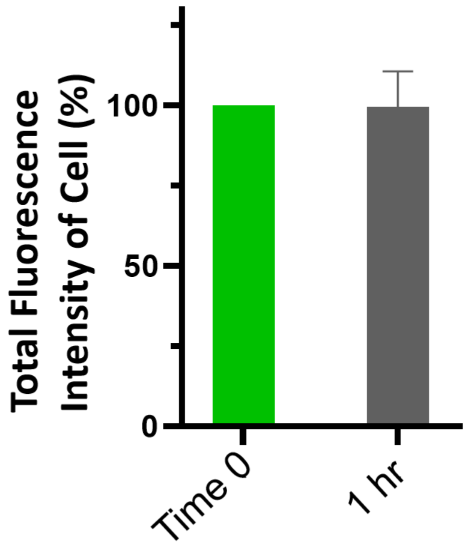


**Fig. S4. Single-level expression levels of PFKL-mEGFP before and after PEG300 treatment.** A total fluorescence intensity of PFKL-mEGFP per cell was quantified from 3D LLSM imaging before and 1-h after PEG300 addition (N = 12). Error bars indicate standard error from at least 3 independent experiments.

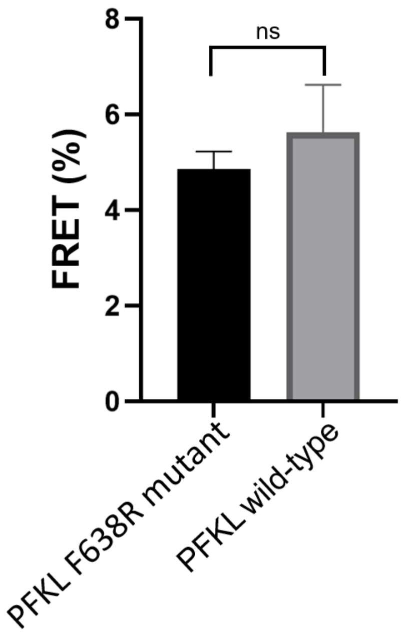


**Fig. S5. FRET measurements from live Hs578T cells.** FRET efficiencies (%) between PFKL F638R mutants and between wild-type PFKLs *outside* glucosomes ($N_{F638R}$ = 48 & $N_{Wild\text{-}Type}$ = 13 ). Error bars indicate standard error from at least 3 independent experiments. Statistical analyses were performed using an unpaired t test. ‘ns’ indicates not significant.

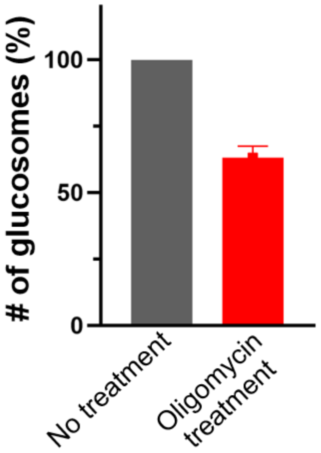


**Fig. S6. Amounts of glucosomes before and after oligomycin A treatment.** We quantified amounts of glucosomes per cell in the absence (grey) and presence (red) of oligomycin A. Error bars indicate standard error from at least 3 independent experiments.

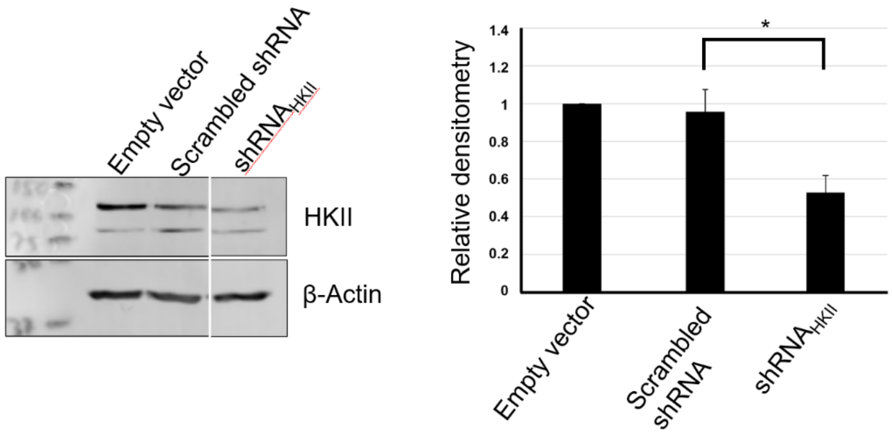


**Fig. S7. Western blot analysis of hexokinase II (HKII) knockdown.** Hs578T cells transfected with empty vector, scrambled shRNA and transfected shRNA$_{HKII}$ were subjected to western blots, showing significant knockdown of HKII in the presence of shRNA$_{HKII}$. β-Actin was used as a load control.

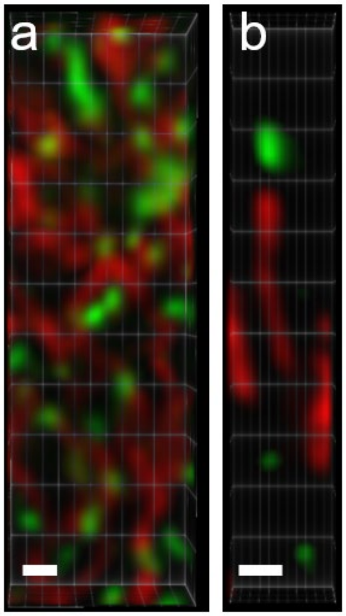


**Fig. S8. Representative images of a mitochondria-dense region (a) and a mitochondria-sparse region (b) from Hs578T cells.** Glucosomes and mitochondria are shown by PFKL-mEGFP (green) and MitoTracker (red), respectively. Scale bars 1 μm.

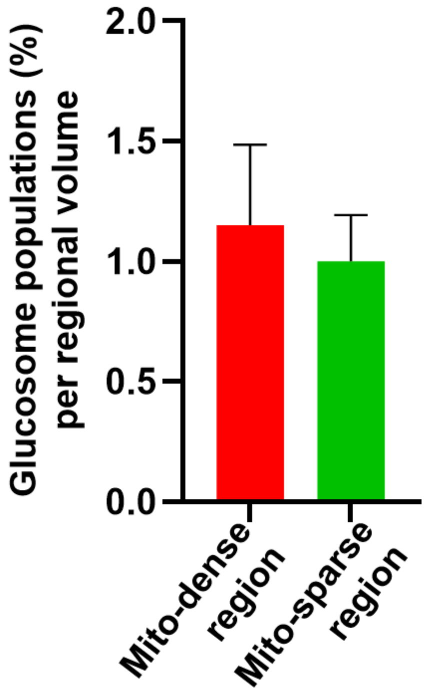


**Fig. S9. Glucosome populations in a mitochondria-dense region (red) and a mitochondria-sparse regions (green) from Hs578T cells.** Error bars indicate standard error from at least 3 independent experiments.

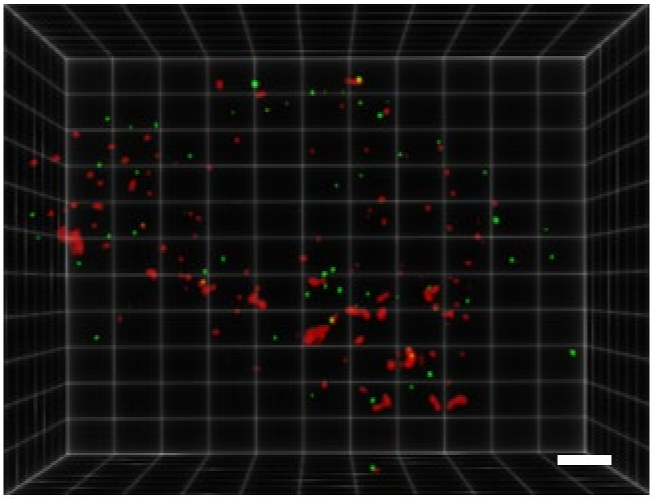

**Fig. S10. A representative Hs578T cell showing co-expression of glucosomes and purinosomes under 3D LLSM imaging.** PFKL-mCherry and FGAMS-mEGFP were used to visualize glucosomes (red) and purinosome (green), respectively. FGAMS, formylglycinamidine ribonucleotide synthase. Scale bars, 5 μm.
